# LARP6 Promotes Features of Plaque Stability and Smooth Muscle Cell Survival in Atherosclerosis

**DOI:** 10.64898/2025.12.22.696106

**Authors:** Meng Zhang, Svitlana Danchuk, Sergiy Sukhanov, Tadashi Yoshida, Patrice Delafontaine, Yusuke Higashi

## Abstract

**Introduction:** Acute plaque rupture or erosion followed by luminal thrombosis is a major cause of myocardial infarction and sudden cardiac death. Vulnerable plaques are characterized by a thin fibrous cap and reduced numbers of smooth muscle cells (SMCs). SMCs are essential for plaque stability, as they synthesize collagen and support fibrous cap structure. Our previous work demonstrated that insulin-like growth factor-1(IGF-1) enhances plaque stability and upregulates La ribonucleoprotein domain family member 6 (Larp6), a collagen mRNA-binding protein in SMCs. However, the specific role of Larp6 in atherosclerotic plaque stability remains undefined.

**Methods:** In vitro, human aortic SMCs (hAoSMCs) were used to investigate the molecular mechanism underlying IGF-1 regulation of LARP6. Using the Myh11 promotor we generated SMC specific Larp6 overexpression mice on an ApoE^-^/^-^ background (SMC-Larp6) to assess effects on plaque stability. Bulk RNA sequencing data from human atherosclerotic plaques (GSE120521) were analyzed to compare stable and vulnerable plaques.

**Results:** IGF-1 regulates LARP6 through phosphorylation and downregulation of microRNAs. Plaques from SMC-Larp6 mice exhibited a significantly higher collagen content, accompanied by a thicker fibrous cap and a smaller necrotic core, increased presence of SMCs, without changes in overall plaque burden compared to controls. LARP6 overexpression in hAoSMCs significantly increased cell proliferation and survival under oxidative stress and promoted collagen accumulation by enhancing collagen synthesis. RNA sequencing analysis of human atherosclerotic plaques revealed reduced expression of LARP6 and downregulation of extracellular matrix-related pathways in unstable plaques, underscoring the clinical relevance of these findings.

**Conclusions:** Larp6 overexpression promotes SMC survival, collagen synthesis and features of plaque stability under atherosclerotic conditions. These findings suggest that Larp6 is a potential therapeutic target for stabilization of atherosclerotic plaques.

## Introduction

Atherosclerotic coronary heart disease (CHO) remains the leading cause of morbidity and mortality worldwide [1]. Most acute coronary events result from the rupture or erosion of atherosclerotic plaques that are often not hemodynamically significant [2]. Therefore, plaque stability is a critical determinant of clinical outcomes.

Stable plaques are characterized by a greater presence of collagen and smooth muscle cells (SMCs) in the fibrous cap, smaller necrotic cores, reduced apoptosis, and fewer inflammatory cells. Collagen constitutes approximately 60% of the total protein content in atherosclerotic plaques[3] and is primarily synthesized by SMCs during the initiation and progression of atherosclerotic lesion formation, as part of the vascular “response-to-injury” mechanism [4, 5]. This process contributes to neointimal thickening. Notably, many acute coronary events occur before the development of occlusive plaques, highlighting the importance of modulating plaque biology to prevent such events [2, 6]. Collagen deficiency contributes to plaque vulnerability and significantly increases the risk of rupture [6, 7]. Thus, collagen content is a key hallmark of plaque stability[4, 8, 9].

Beyond quantity, collagen quality also plays a crucial role. Type I collagen forms a complex fibrillar structure, and macrophages cultured on polymerized collagen secrete lower levels of MMP9 [1O], thereby reducing collagen degradation. Efficient translational or post-translational regulation of collagen assembly likely promotes the formation of an extracellular matrix that provides an anti-atherogenic microenvironment for plaque-resident cells[8, 11].

LARP6/Acheron, first identified in Tobacco hawk moth *(Manduca sexta)* in 2007 [12], regulates muscle development and cell death [13–15]. Its mammalian ortholog, LARP6 (La ribonucleoprotein domain family member 6), is an RNA-binding protein that interacts with the mRNAs encoding type I (COL1A1 and COL1A2) and type Ill (COL3A1) collagen, regulating their translation and promoting fibril formation [16]. LARP6 binds a conserved stem-loop structure in the 5’-untranslated regions (5’-UTRs) of these mRNAs. Interestingly, mutations that disrupt this stem-loop motif reduce collagen production and result in pepsin-sensitive collagen deposition, indicating defective fibril assembly [17].

We previously demonstrated that insulin-like growth factor 1 (IGF1) reduces atherosclerotic burden and promotes plaque stability in Apoe-^1^- mice [18, 19], potentially through enhanced collagen matrix deposition and maturation mediated by the upregulation of LARP6 [20]. In the present study, we further investigated the molecular mechanisms by which IGF-1 regulates LARP6 expression and activity. Building on these findings, we hypothesized that SMC-specific overexpression of LARP6 would enhance plaque stability via increased collagen synthesis and improved fibrillar organization., In this study, we have developed a smooth muscle cell-specific Larp6 gain-of-function mouse model (Larp6sMc-oE) to test this hypothesis.

## Materials and Methods

### Animals

All animal procedures were conducted in accordance with protocols approved by the Tulane University Institutional Animal Care and Use Committee. Mice were housed in a temperature-controlled room (22 °C) under a 12-hour light/12-hour dark cycle and provided ad libitum access to water and a diet. For atherosclerosis study, 8 weeks old animals after tamoxifen administration (see below) were fed on a Western diet (Envigo TD.88137), containing 42% fat, 34% carbohydrates, and 2% cholesterol, for 12 weeks before terminal tissue collection.

### Bolus IGF-1 administration

Recombinant human IGF-1 (Ipsen Biopharmaceuticals, 10.mg/mL) was administered via intraperitoneal (IP) injection 18 hours prior to euthanasia. A dose of 1.5 mg/kg body weight was used, delivered in a 100µL injection volume for mice weighing 20-35 g. The drug was aseptically diluted with pharmaceutical-grade saline to achieve the appropriate concentration.

### Oil Red O Assay

Mice were euthanized under deep anesthesia with an intraperitoneal injection of ketamine (100 mg/kg) and xylazine (12 mg/kg), dosed according to body weight. Following anesthesia, the animals were perfused via the left ventricle as previously described[21]. The entire aorta was then dissected and fixed overnight in formaldehyde. After removing adventitial fat, the aorta was stained with Oil Red 0, opened longitudinally, pinned en face, and imaged using a Leica stereomicroscope. Total arterial surface area and lesion area were quantified using lmageJ. Atherosclerotic burden was expressed as the percentage of the total arterial area covered by Oil Red O-positive lipid-rich lesions.

### Plaque composition

Mice were anesthetized and perfused transcardially with saline followed by 4% paraformaldehyde (buffered) containing 5% sucrose. Hearts were dissected, post-fixed overnight, and embedded in paraffin. Serial 5 µm sections were collected throughout the entire aortic valve area (AVA) and stained with Masson’s Trichrome for histological analysis.

To quantify plaque morphology, two representative sections per AVA were imaged using a slide scanner (ZEISS Axio Scan.Z1). Lesion area, necrotic core size, and collagen content were measured using cellSens software (Olympus). Fibrous cap thickness was assessed using a custom lmageJ macro. One hundred measurements were taken between the top of the necrotic core and the luminal edge of the plaque, and the minimum value was recorded as the cap thickness.

### lmmunofluorescence

Serial 5 µm paraffin-embedded cross-sections were collected throughout the entire aortic valve area (AVA). Two sets of serial sections were used to measure smooth muscle cell (SMC) and macrophage areas. Sections were incubated with Alexa Fluor 674-conjugated anti-a-smooth muscle actin antibody (Abeam, 1:200) or rat anti-mouse Mac-3 monoclonal antibody (BioLegend, 1:20), with isotype-matched rat lgG (Abeam) as a negative control. Detection was performed using a biotinylated secondary antibody and an avidin-peroxidase complex (Vectastain Elite ABC Kit, Vector Laboratories).

Signals were developed with a DAB substrate kit (lnvitrogen), and nuclei were counterstained with DAPI (lnvitrogen). Apoptosis was assessed by TUNEL assay (AssayGenie) according to the manufacturer’s protocol. Briefly, paraffin sections were dewaxed for 30 min, treated with proteinase K for 20 min, and incubated with TdT labeling solution for 60 min at 37 °C.

### Cell culture

Human aortic smooth muscle cells (HASMCs; Lonza) were cultured in Smooth Muscle Basal Medium (SmBM; Lonza) supplemented with 5% fetal calf serum (FCS), antibiotics, recombinant human epidermal growth factor, insulin, and recombinant human fibroblast growth factor. Experiments were conducted using cells at passages 4-8 under serum-free conditions in a 1:1 mixture of Dulbecco’s Modified Eagle Medium (DMEM) and F-12 nutrient solution (lnvitrogen). Mouse vascular smooth muscle cells (MOVAS; ATCC) were cultured in DMEM (Gibco) supplemented with 10% fetal bovine serum (FBS) at 37 °C in a humidified incubator with 5% CO_2_.

For LARP6 overexpression, cells were transduced with adenoviral vectors (Vector BioLabs, Philadelphia, PA) at a multiplicity of infection (MOI) of 40 for 24 hours. Plasmids encoding wild-type mouse Larp6 and phospho-mutant SA-Larp6 were kindly provided by Dr. Branko Stefanovic. Transient transfections were performed using 2.5 µg of plasmid DNA and Lipofectamine 3000 (Thermo Fisher Scientific), with a 48-hour incubation. To assess Larp6 protein stability, cells were treated with cycloheximide (100 µg/mL; Thermo Scientific) following adenoviral transduction.

For cell survival assays, oxidized low-density lipoprotein (oxLDL; Kalen Biomedical LLC) was applied at 100 µg/mL to induce apoptosis. Apoptotic cells were detected using Annexin V (1:200 dilution; Sartorius) in the presence of 1 mM CaCl_2_, according to the manufacturer’s instructions. Real-time cell imaging was performed using the lncucyte Live-Cell Analysis System (Sartorius) under both brightfield and fluorescence channels over a 24-hour period.

### Hydroxyproline assay

Hydroxyproline content was quantified using the Hydroxyproline Assay Kit (Sigma-Aldrich, MAK463) according to the manufacturer’s instructions. Protein lysates from cells or animal tissues were first analyzed for total protein concentration using the BCA assay (Thermo Scientific). For hydroxyproline measurement, 30 µg of tissue lysate or 10 µg of cell lysate protein was used per reaction.

### Western Blot

Western blot analysis was performed as previously described[21]. Briefly, cells or animal tissues were washed with cold phosphate-buffered saline (PBS) and lysed in radioimmunoprecipitation assay (RIPA) buffer containing 150 mM NaCl, 50 mM Tris-HCI (pH 8.0), 1% Triton X-100, 0.5% sodium deoxycholate, and 0.1% SOS. Animal tissues were homogenized using a Bullet Blender (Next Advance). Protein concentrations were determined using the bicinchoninic acid (BCA) protein assay (Thermo Scientific). Equal amounts of protein were mixed with loading buffer containing 13-mercaptoethanol, resolved by 10% SOS-PAGE, and transferred to membranes for immunoblotting. The following primary polyclonal antibodies were used: collagen type I (Rockland, 1:1000), LARP6 (Thermo Fisher Scientific, 1:1000), phospho-Akt (Thr308; Cell Signaling Technology, 1:1000), collagen I a1 C-telopeptide (Rockland, 1:1000), and 13-actin (Sigma, 1:20,000) as a loading control. lmmunoreactive bands were detected using enhanced chemiluminescence (ECL; Thermo Fisher Scientific), imaged with a GelDoc system (Bio-Rad), and quantified using Image Lab software (Bio-Rad).

### MicroRNA Isolation and Quantification

Human aortic smooth muscle cells (hAoSMCs) were cultured and exposed to IGF-1 (100 ng/ml) for 3 hours. Total RNA, including small RNAs, was isolated using the miRNeasy Mini Kit (Qiagen) according to the manufacturer’s protocol. The purity and concentration of RNA were assessed spectrophotometrically. cDNA was synthesized using the miScript II RT Kit (Qiagen) with the HiSpec buffer to ensure selective reverse transcription of mature miRNAs. qRT-PCR was performed using the miScript SYBR Green PCR Kit (Qiagen) and specific miScript primer assays for target miRNAs. Expression levels of each miRNA were normalized to SNORD61.

### Bulk RNA seq analysis

Publicly available RNA-seq data were obtained from the NCBI Gene Expression Omnibus (GEO; accession number GSE120521). Data analysis was performed using iDEP v2.1 (bioinformatics.sdstate.edu/idep). Raw read counts were uploaded, filtered to exclude lowly expressed genes, and normalized using variance-stabilizing transformation (rlog). Sample relationships were explored by principal component analysis (PCA) on rlog-transformed data to assess variance among experimental groups. Differential expression analysis was carried out with the DESeq2 pipeline integrated into iDEP. Genes meeting the significance criteria (adjusted p-value < 0.05, Benjamini-Hochberg correction) were considered differentially expressed. Lists of differentially expressed genes were further analyzed for functional enrichment within iDEP. Overrepresentation of biological pathways was assessed using KEGG and Reactome databases, with expressed genes serving as the background. Fisher’s exact test was used to calculate enrichment significance, and multiple testing correction was applied.

### Statistical analysis

All numerical data are presented as mean± SEM. Statistical analyses were performed using GraphPad Prism software (GraphPad Software). Data distributions were first evaluated with the D’Agostino-Pearson omnibus normality test, and equality of variances was assessed using the F test. Between-group comparisons were performed using an unpaired Student’s *t* test or the Mann-Whitney U test, depending on the normality of the residuals. A two-tailed *P* value < 0.05 was considered statistically significant.

## Result

### IGF-1 induces phosphorylation of LARP6

Our previous work demonstrated that IGF-1 promotes collagen synthesis via upregulating Larp6 expression[21]. To validate this mechanism in vivo, we administered IGF-1 to mice and assessed collagen synthesis by quantifying hydroxyproline levels in aortas. Consistently, IGF-1significantly increased collagen synthesis in mouse aortic protein lysates (Figure 1A). Furthermore, Larp6 overexpression in SMC similarly enhanced collagen production compared to controls. Notably, IGF-1 administration in the setting of Larp6 overexpression did not further increase hydroxyproline levels compared to saline-treated mice, suggesting that the effect of IGF-1 on collagen deposition is largely dependent on Larp6.

**Fig. 1.**
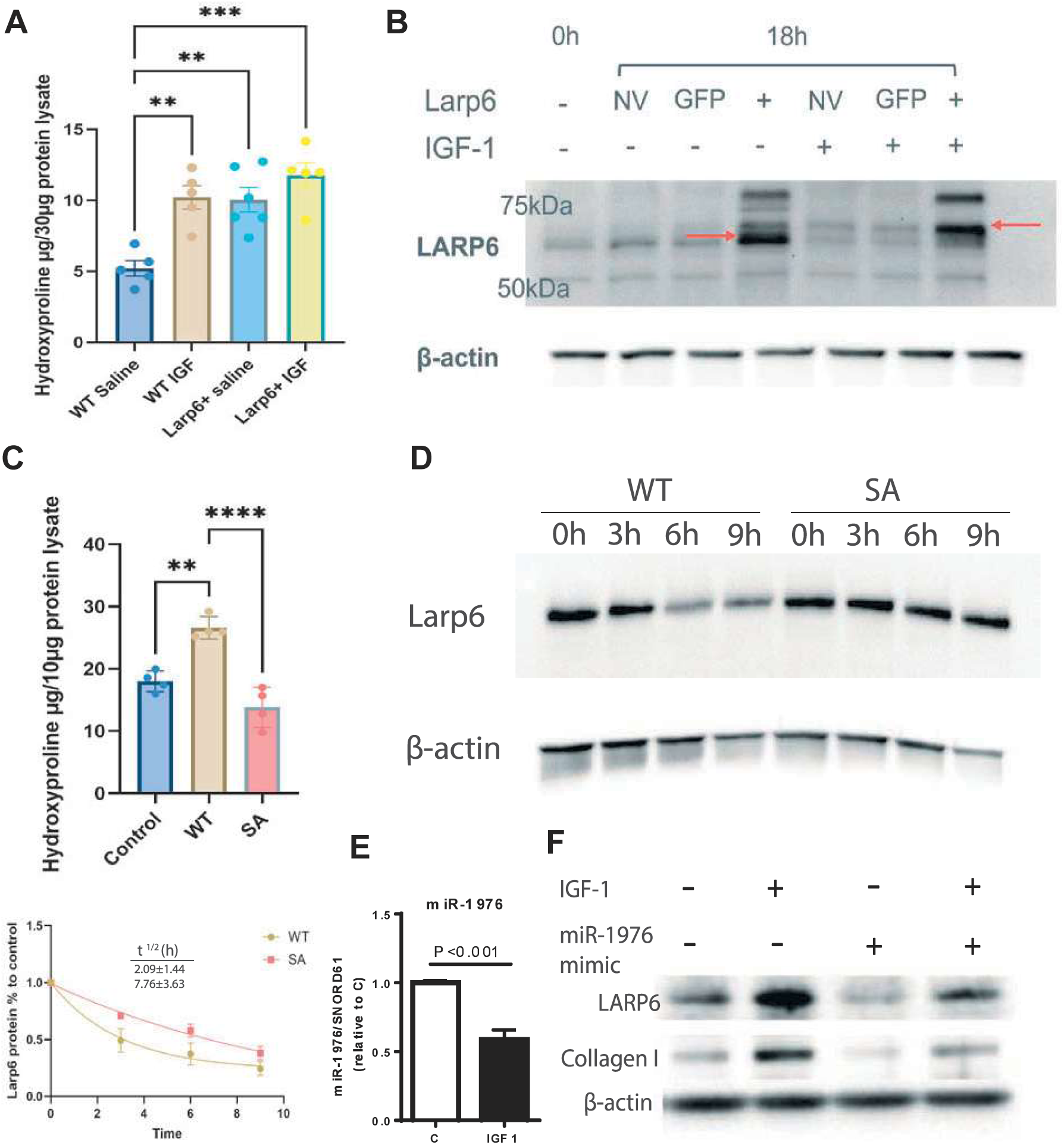
Larp6 is regulated by IGF-1 via phosphorylation and micro-RNAdependent mechanisms. **A,** Collagen production in mouse aortas measured by hydroxyproline assay. WT or Larp6^SMC-OE^ mice were injected with IGF-1 (1.5mg/kg-BW, i.p.), and aortas were dissected 18h later and homogenized to measure hydroxyproline contents. Data are presented as mean ± SEM.; statistical analysis was performed using one-way ANOVA(**P <0.01, ***P< 0.001). **B,** Western blot analysis detecting LARP6 protein in human aortic smooth muscle cells (hAoSMCs) (*n* = 3 per group). Cells were transduced with adenovirus vector for 24 hours, using no-virus (NV) and GFP controls, serum-starved overnight, and then treated with IGF-1 (50 µg/mL) for 18 hours. **C,** Collagen production in MOVAS cells assessed by hydroxyproline assay after 24-hour adenoviral overexpression of either WT or SA mutant Larp6, followed by an additional 18-hour incubation. **D,** Larp6 protein stability in MOVAS cells transfected with plasmids encoding WTor SA mutant Larp6, followed by cycloheximide treatment for 9 hours. One-phase decay models were fitted, and half-lives were calculated from three independent experiments (*n* = 5 per group). **E,** Quantitative RT-PCR analysis of miR-1976 expression in hAoSMCs following IGF-1 treatment. Expression levels were normalized to SNORD61 and presented relative to control. **F,** Western blot analysis of LARP6 and collagen type I (Col1) protein levels in hAoSMCs treated with a miR-1976 mimic for 24 hours, followed by 3 days of serum starvation and then 4 days of IGF-1 treatment.

To elucidate how IGF-1 regulates LARP6, we revisited our previous finding that IGF-1 does not alter Larp6 mRNA levels [21], suggesting a post-transcriptional regulation. Western blot analysis of IGF-1-treated hAoSMCs revealed two distinct LARP6 protein bands at 60kDa and 62kDa (Figure 1B). Upon IGF-1 stimulation, a clear shift from the lower to the upper band was observed, consistent with a post-translational modification (PTM). This band shift was evident for both endogenous and adenovirally overexpressed LARP6 (Figure 1B).

We interrogated phosphoproteomic databases and identified several phosphorylation sites on LARP6. In contrast, only a single acetylation and a single methylation site were detected at amino acids 2 and 41, respectively (Figure S1A, Table S1). Given that acetylation and methylation are unlikely to produce detectable shifts in SOS-PAGE mobility, phosphorylation emerges as the most plausible PTM responsible for the observed band shift in response to IGF-1. Consistent with this hypothesis, our previous work showed that inhibition of Pl3K and AKT abrogated both the IGF-1-induced band shift and the concomitant increase in collagen synthesis[21]. Extending our prior findings, which identified AKT as the upstream kinase, Zhang et al. identified serine 451 (S451) as the specific AKT-mediated phosphorylation site on LARP6 [22].

To assess the functional significance of S451 phosphorylation in smooth muscle cells, we generated a phospho-null S451A mutant of mouse Larp6 and transfected it in mouse aortic smooth muscle cells (MOVAS). While wild-type Larp6 significantly increased hydroxyproline levels, the S451A mutant exhibited a markedly reduced capacity to increase collagen synthesis (Figure 1C). These results indicate that phosphorylation at S451 is essential for LARP6-mediated promotion of collagen synthesis.

We further investigated whether S451 phosphorylation influences LARP6 protein stability. MOVAS cells overexpressing either wild-type or S451A mutant Larp6 were treated with cycloheximide to inhibit new protein synthesis. Surprisingly, the S451A mutant exhibited a longer half-life than wild-type Larp6 (Figure 1D), suggesting that phosphorylation at S451 promotes LARP6 turnover. Collectively, these findings suggest that AKT-mediated phosphorylation at S451 is critical not only for LARP6’s collagen-promoting function but also for the regulation of its protein stability in smooth muscle cells.

### IGF-1 regulates LARP6 protein levels by downregulating microRNAs

Although our previous work demonstrated that IGF-1 increases LARP6 protein levels [21], the underlying mechanism remained unclear. Given thatAKT-mediated phosphorylation at S451 promotes LARP6 degradation, phosphorylation alone cannot account for the observed increase in protein abundance. This discrepancy prompted us to investigate potential alternative mechanisms whereby IGF-1 elevates LARP6 levels..

MicroRNAs (miRs) are well-established post-transcriptional regulators of gene expression and are implicated in cardiovascular pathophysiology. Through database mining, we identified six candidate miRs that target LARP6 and are downregulated by IGF-1. All six miRs were expressed in hAoSMCs, and among them, miR-126-3p, miR-744-5p, and miR-1976 show significant suppression upon IGF-1 treatment (Figure 1E, Table S2). Functional assays demonstrated that miR-1976 markedly reduced LARP6 protein levels, an effect that was attenuated by IGF-1 treatment (Figure 1E). Taken together, these results suggest that IGF-1 regulates LARP6 through two complementary mechanisms: phosphorylation, which modulates its activity and turnover, and miR-mediated repression, which controls its protein abundance.

### A SMMHC-CreERT^2^ Mouse Model for Smooth Muscle Cell Conditional Larp6 Overexpression

Given that the regulation of collagen synthesis by IGF-1 is largely dependent on Larp6we hypothesized that Larp6 overexpression in SMC would enhance collagen deposition and promote plaque stability under atherosclerotic conditions. To test this hypothesis, we generated SMC-specific tamoxifen-inducible Larp6 overexpressing mice by crossing SMMHC-CreERT2 (Jackson Lab: Stock#019079) and Tg(CAG-CAT-Larp6) (“CAG-CAT-Larp6”); both on Apoe-deficiency background. Following tamoxifen administration, Cre recombinase is activated under the control of the SM-MHC promoter, excising the CAT coding sequence and its stop codon, thereby enabling Larp6 expression under the control of the CAG promoter in SMCs.

To induce Larp6 overexpression, 8-week-old Larp6sMc-oE mice and SMMHC-CreERT2 littermate controls received tamoxifen. All mice were then fed a Western diet (WO) for 12 weeks (Figure 2A). Western blotting confirmed markedly elevated Larp6 protein levels in SMC-enriched tissues, such as aorta and urinary bladder, consistent with successful transgene activation (Figure 2B). Plasma cholesterol levels did not differ between Larp6SMC-OE mice and control mice (Figure 2C).

**Fig. 2.**
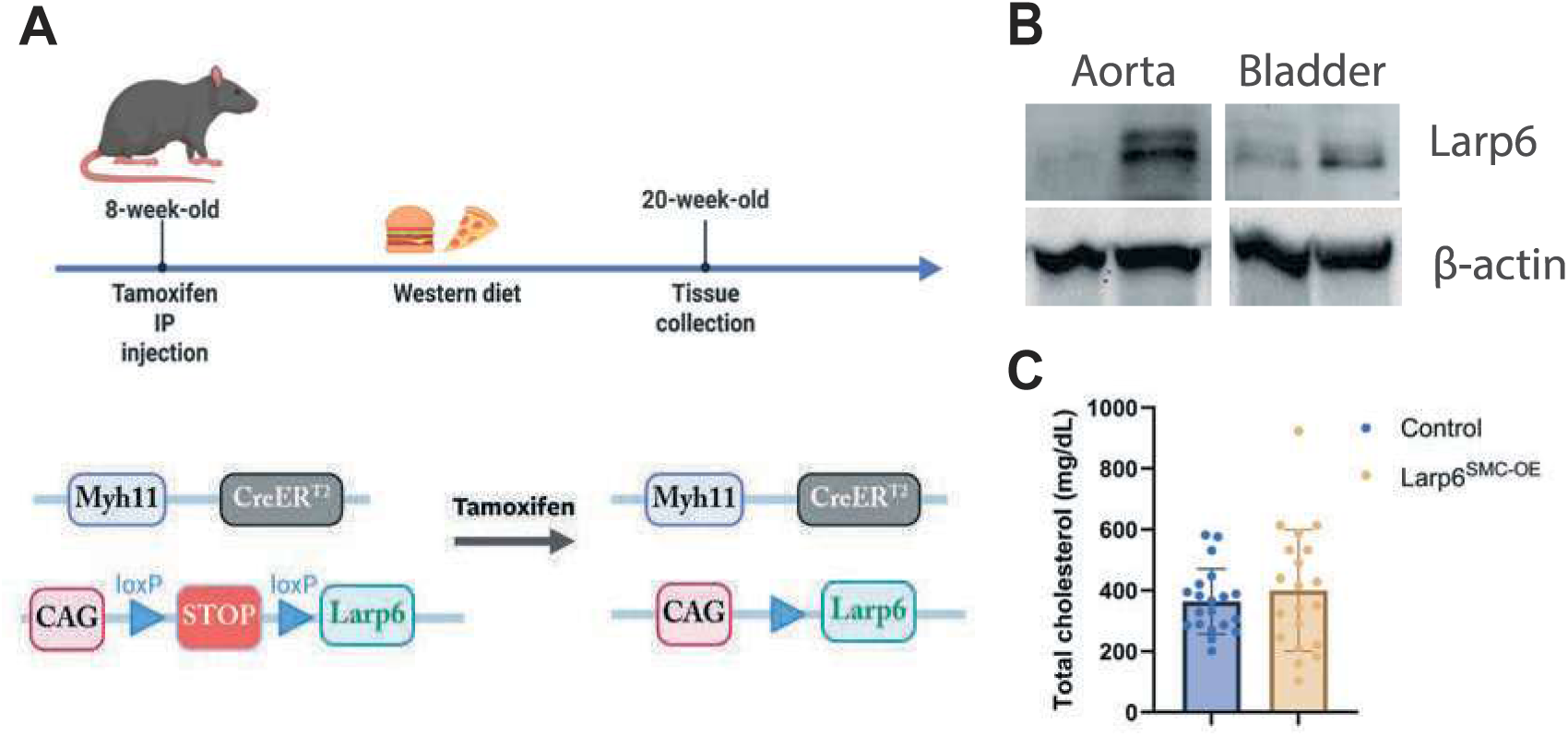
Overexpression of Larp6 in Larp6S^MC-OE^ mice. **A,** Larp6 overexpression was induced via tamoxifen administration (injected i.p. at 75 mg/kgBW/day for 5 consecutive days). Following induction, mice were fed a Western diet(WD) for 12 weeks. Only male mice were used, as the *Myh11-CreERT2* transgene is integrated into the Ychromosome. **B,** Larp6 protein expression was elevated in Larp6^SMC-OE^ mice across SMC-enriched organs. **C,** Serum cholesterol levels after 12-week WD feeding (n=20 mice per group). Data are mean ±SEM. Graphic in **A** was created with BioRender.

### SMC specific Larp6 overexpression increases collagen content in mouse plaque without changing plaque burden

To assess atherosclerotic burden, we stained the entire aorta from Larp6sMc-oE and control with Oil Red 0. *En face* analysis demonstrated no significant difference in the plaque-covered area between groups (Figure 3A). To evaluate plaque collagen content and composition in Larp6SMC-OE mice, we performed Masson’s Trichrome and Picrosirius Red staining on aortic root sections (Figure 3B-G). Consistent with *en face* analysis, plaque size in the aortic root did not differ significantly between Larp6SMC-OE and control mice. However, Larp6sMc-oE plaques exhibited markedly increased collagen levels, accompanied by a thicker fibrous cap and a smaller necrotic core-features indicative of enhanced plaque stability.

**Fig. 3.**
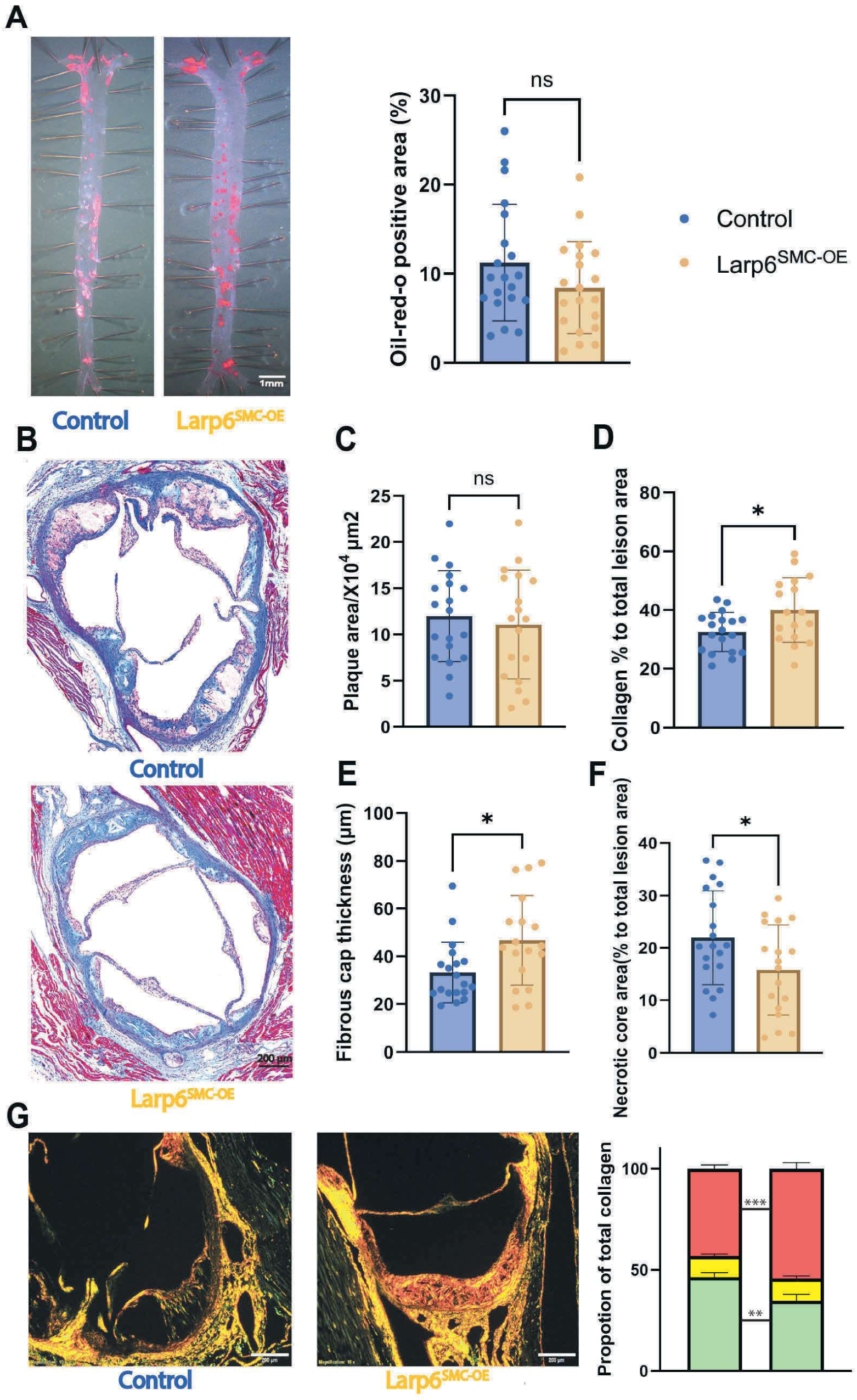
Larp6^SMC-OE^ increases collagen content in mouse plaque without altering the plaque burden. **A,** En face Oil Red O staining of whole aortas from Larp6^SMC-OE^ and littermate control mice to assess plaque area (*n* = 20 per group). Data are presented as mean ± SEM. Scale bar = 1mm **B-F,** Masson’s Trichrome staining to determine collagen contents within atherosclerotic lesions. B, Representative images showing total collagen deposition (blue) within lesions. Scale bar = 200µm. **C,** Quantification of the lesion size and **D,** collagen-content. **E,** Quantification of the fibrous cap thickness at the thinnest region and **F,** necrotic core size. Data are presented as mean ± SEM. \*\**P*< 0.01 by Student’s *t*-test. **G,** Picrosirius Red staining under polarized microscopy, red, green, yellow colors are compared by Student’s t-test, ± SEM, \*\**P*<0.01, \*\*\**P*<0.001.

We validated the collagen increase in Larp6SMC-OE mice by Picrosirius Red staining, which identifies fibrillar collagen networks in tissue sections (Figure S2A). During atherosclerosis progression, elevated levels of matrix metalloproteinases contribute to collagen degradation, thereby weakening the fibrous cap [23]. Under polarized microscopy, the color of Picrosirius Red stained collagen reflects fiber thickness and organization: thinner, loosely packed fibers appear green, whilst thicker, well-organized fibers shift toward yellow to red birefringence [24]. In Larp6SMC-OE mice, collagen fibers showed a marked increase in red birefringence, indicating enhanced thickness and structural integrity (Figure 3G).

Together, these findings suggest that Larp6 overexpression in SMCs promotes a more stable plaque phenotype by reinforcing the collagen matrix, without altering overall plaque burden.

### Larp6 overexpression increases SMC presence in the plaque

Stable atherosclerotic plaques are characterized by a thicker fibrous cap and increased smooth muscle cell content. To examine whether Larp6 influences SMC presence within plaques, we performed immunofluorescence staining for a-smooth muscle actin (a-SMA) in aortic root sections (Figure 4A). Larp6sMc-oE mice exhibited a significant 80.0 ± 26.9% increase in SMC content compared with controls. As inflammation and cytokine activity contribute to the vulnerability of plaque[25], we also assessed macrophage abundance within the lesions and found no significant difference between Larp6sMc-oE and the control plaques (Figure S2B). Given prior studies linking LARP6 to enhanced cell survival[26] and proliferation[27], we evaluated SMC apoptosis within the plaques using TUNEL staining co-labeled with a-SMA. Larp6sMc-oE mice displayed a significant reduction in apoptotic SMCs compared with controls (Figure 4B).

**Fig. 4.**
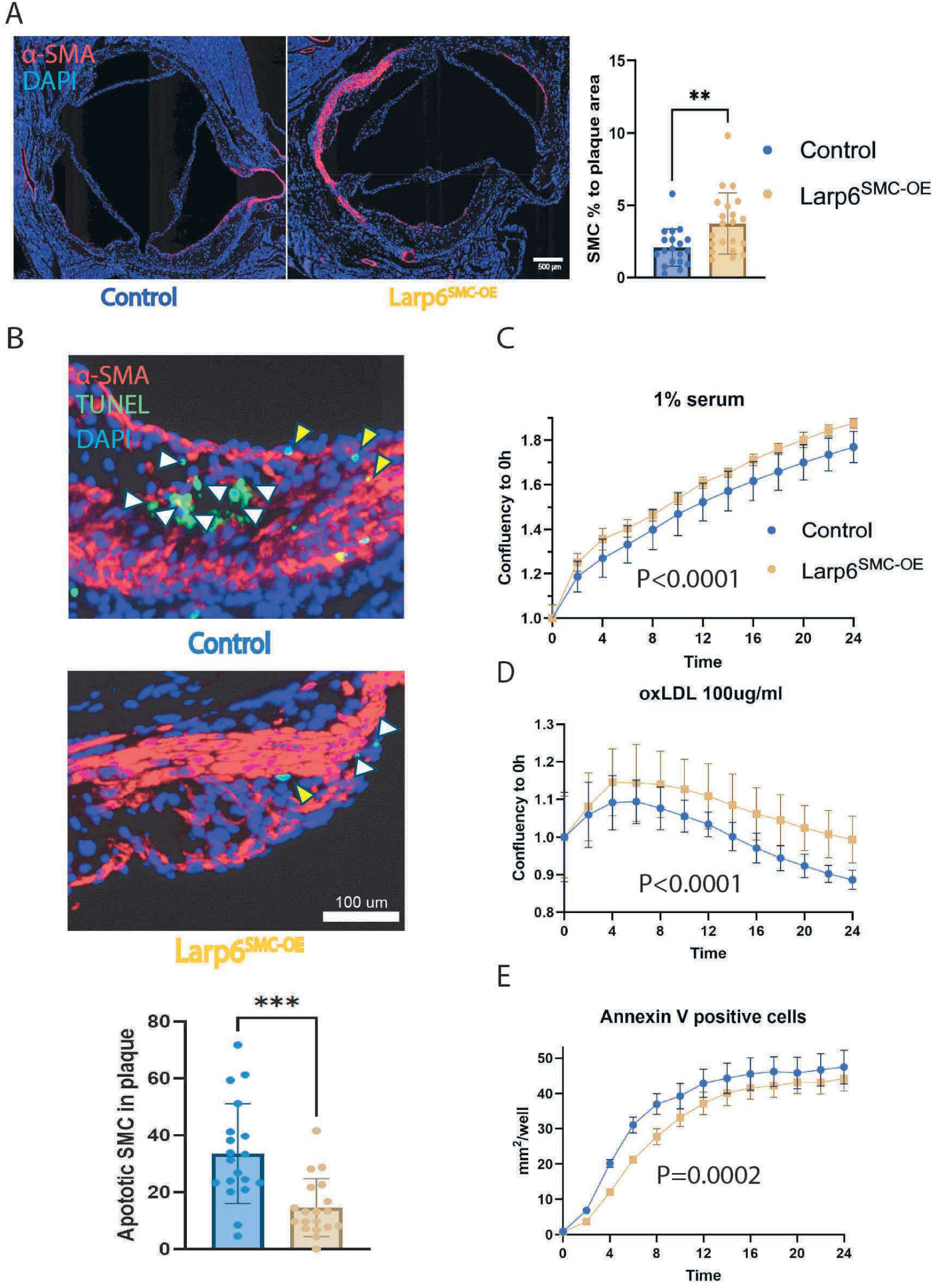
A, Larp6 OE increased SMC presence in the plaque. **A,** Representative images for α-smooth muscle actin (αSMA) staining (red) counterstained with DAPI (blue), showing smooth muscle cell distribution in plaque from control or Larp6^SMC-OE^ mice after 12-week of WD feeding. Quantification of αSMA-positive area is expressed as a percentage of total plaque area. Data are presented as mean ± SEM. \*\**P*< 0.01 by Student’s *t*-test. **B,** Representative immunofluorescence images of aortic root plaques from control and Larp6^SMC-OE^. Yellow arrowheads indicate apoptotic SMCs (TUNEL^+^α-SMA^+^), whereas white arrowheads denote apoptotic non-SMC cell types. Scale bar = 100 µm. **C - E,** Human aortic smooth muscle cells (HASMCs) were seeded at 5,000 cells per well in 96-well plates and treated under various conditions to assess proliferation (**C** and **D**) and apoptosis (**E**). **C,** Cells were cultured under 1% serum condition or **D,** 100µg/ml oxidated LDL (oxLDL) and cell confluency was monitored overtime, normalized to baseline (0 h), and exponential growth curves were fitted for analysis. **E,** Cells were exposed to 100 µg/mL oxLDL along with Annexin V-FITC to detect apoptosis over a 24-hour period. Annexin V-positive cell-covered area was measured and compared. Data represents mean ± SEM from four biological replicates.

To further elucidate underlying mechanisms, we used human aortic SMCs (hAoSMCs) and overexpressed LARP6 to investigate the role of LARP6 in cell proliferation and survival. Cells were transduced with adenovirus to overexpress LARP6, which significantly enhanced SMC proliferation under 1% serum conditions (Figure 4C). To assess cell survival under conditions simulating atherogenic stress, cells were exposed to increasing concentrations of oxidized LDL (oxLDL), with native LDL (nLDL) serving as a control. OxLDL doses 2:50 µg/ml induced substantial cell death (Figure 4D). Notably, at 100 µg/ml oxLDL, LARP6-overexpressing cells exhibited significantly higher viability than controls after 24 hours (Figure 4D). Consistently, apoptosis analysis using Annexin V staining demonstrated that LARP6 overexpression significantly decreased the number of apoptotic SMCs over a 24-hour period (Figure 4E). These data collectively suggest that LARP6 promotes both the proliferation and survival of SMCs under atherosclerotic conditions, suggesting that LARP6 contributes to the development of more stable, SMC-enriched plaques.

### LARP6 expression is decreased in human unstable atherosclerotic plaque

To obtain further insights into the potential role of LARP6 in human atherosclerosis, we analyzed publicly available RNA-seq data from patient-derived atherosclerotic plaques (GSE120521)[28], which had been spatially classified into stable and unstable regions. Heatmap visualization and principal component analysis (PCA) demonstrated a clear separation between these two plaque phenotypes (Figure S3A-B). Differential expression analysis identified 1,568 genes significantly upregulated and 1,349 genes downregulated in unstable plaques. Notably, LARP6 expression was significantly reduced in unstable regions (Figure 5A-B).

**Fig. 5.**
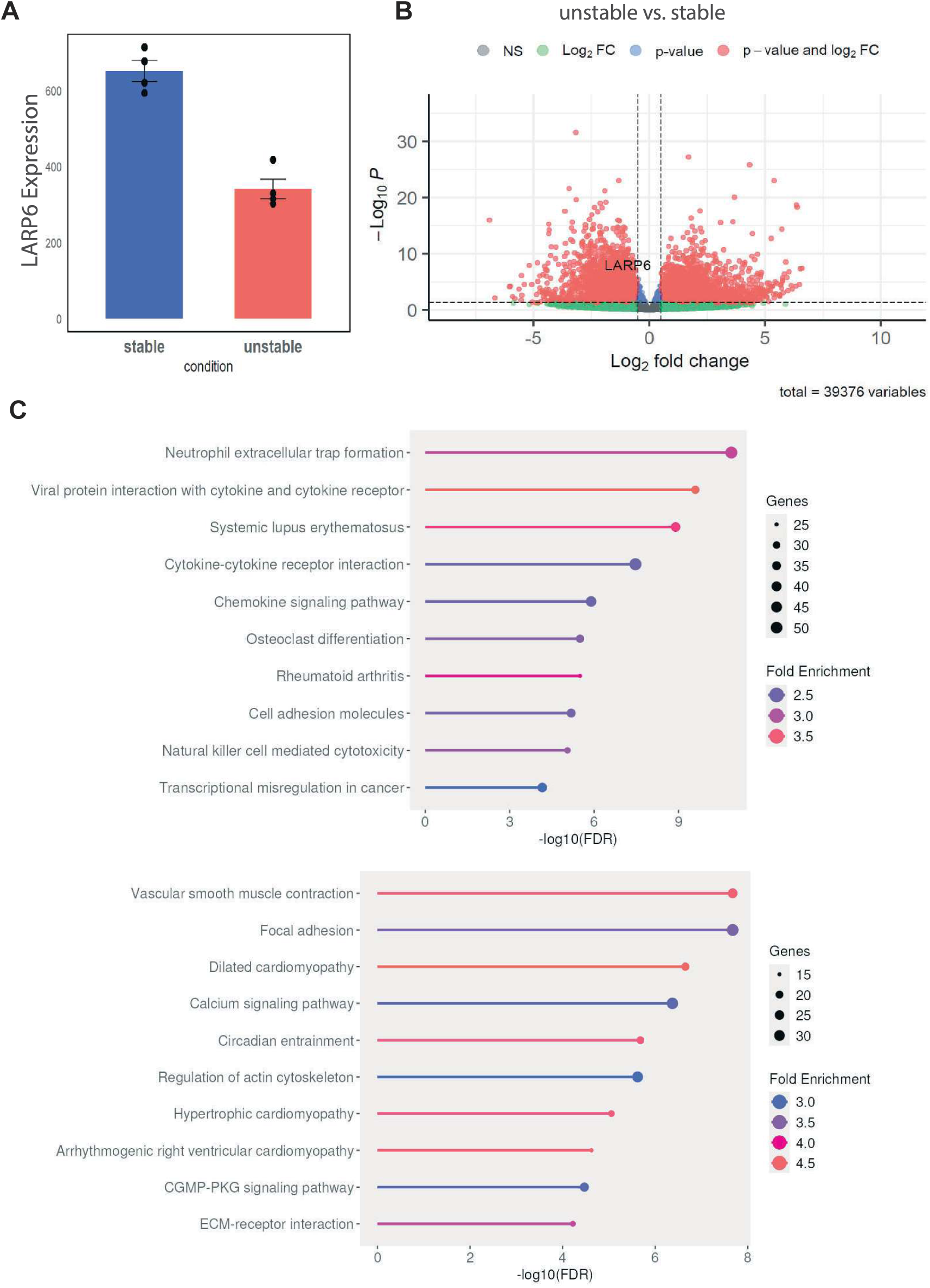
Larp6 decreased in human unstable plaque. **A,** LARP6 expression levels in human atherosclerotic plaques, comparing stable and unstable lesions and **B,** DEGs identified by bulk RNA-sequencing, analyzed using the Wald test with DESeq2. **C,** Pathway enrichment analysis comparing unstable versus stable plaques, with significant pathways identified using Gene Set Enrichment Analysis.

To gain insight into the pathways associated with plaque instability, we performed KEGG enrichment pathway analysis on the differentially expressed genes. Inflammatory pathways, including cytokine-cytokine receptor interaction, chemokine signaling, and osteoclast differentiation, were prominently upregulated in unstable plaques (Figure 5C). In contrast, pathways related to structural integrity and extracellular matrix (ECM) maintenance, such as smooth muscle contraction, focal adhesion, and ECM-receptor interaction, were downregulated. Notably, focal adhesion and ECM-receptor interaction are directly involved in collagen biosynthesis, crosslinking, and matrix anchoring-highlighting a potential mechanistic link between reduced LARP6 expression and impaired plaque stability. Together, these results suggest that LARP6 is downregulated in unstable human plaques, and that its role in collagen regulation may represent a critical component of the molecular program underlying plaque stabilization.

## Discussion

The pathophysiological hallmarks of atherosclerotic plaque instability include intraplaque hemorrhage[29], excessive lipid accumulation[30], inflammation, increased cell death and the presence of a thin fibrous cap[31]. In recent years, extensive research has focused on identifying molecular pathways underlying plaque instability with the aim of developing new therapeutic strategies. These strategies include modulation of lipid metabolism to reduce cholesterol burden[32], regulation of programmed cell death and efferocytosis[33], attenuation of inflammatory and immune responses[34] and reduction of oxidative stress through control of reactive oxygen species[35]. Although each mechanism can function independently, they are highly interconnected, forming a complex regulatory network that determines plaque vulnerability. Within this complex interplay, the ECM serves as the structural backbone of the plaque. A well-organized ECM, rich in fibrillar collagen, provides tensile strength to the fibrous cap, limits rupture, and maintains local microenvironmental homeostasis.

In this study, we provide the first evidence implicating LARP6 in the regulation of plaque stability and SMC behavior during atherogenesis. Overexpression of Larp6 in SMCs led to a marked increase in plaque collagen content, contributing to enhanced fibrous cap formation and a reduction in necrotic core size, consistent with overall plaque stabilization. We also observed a significant increase in SMC content in Larp6^5^MC-OE plaques. SMCs are widely recognized as beneficial for plaque stability, as SMCs synthesize ECM components that reinforce structural integrity. However, during disease progression, SMCs can undergo phenotypic modulation-losing contractile markers and adopting fibroblast-, macrophage-, or osteochondrocyte-like phenotypes[36]. These phenotypically altered SMCs might become foam cell-like and/or proinflammatory, secreting cytokines that accelerate lesion progression. Using a-smooth muscle actin as a marker of contractile SMCs, we identified an enrichment of SMCs in Larp6^5^MC-oE plaques, suggesting a more stable, less inflammatory plaque phenotype. Although macrophage content did not differ significantly between groups, our findings indicate that LARP6 may promote a plaque environment resistant to inflammatory remodeling. Further studies will be required to define the inflammatory landscape and molecular mediators downstream of LARP6 within the plaque microenvironment.

In our previous study, we demonstrated that systemic IGF-1 infusion increases features of plaque stability in Apoe-deficient mice, [21]. Pioneering work by Stefanovic and colleagues has extensively characterized LARP6’s role in collagen regulation in the setting of fibrotic diseases. Using cultured fibroblasts , they demonstrated that LARP6 specifically binds to a conserved 5’ stem-loop structure present in COL1A1 and COL1A2 mRNAs[37], thereby stabilizing collagen transcripts and enhancing their translational efficiency. In the current study, we further dissected the molecular mechanisms underlying IGF-1 regulation of LARP6. Our findings reveal two major regulatory pathways: phosphorylation-dependent activation of LARP6, and microRNA-mediated suppression of its protein levels. Phosphorylated LARP6 induced more collagen synthesis (Figure 1C). Interestingly, phosphorylated LARP6 exhibited reduced stability compared to its phosphorylation-null mutant, suggesting a higher turnover rate of the activated form. Together, these findings suggest that IGF-1 enhances both the activation and synthesis of LARP6, leading to sustained upregulation of collagen production. [38]

Our data suggested that LARP6 promotes cell survival (Figure 4B, E). The ortholog of Larp6, known as Acheron, was first described by Sheel, A., et al. [26] , where it was identified as a gene associated with programmed cell death during development of muscle tissue in hawkmoths [26]. Subsequent studies demonstrated that Acheron binds to a novel BH3-only protein, BAD/BNIP3-homolog, thereby inhibiting apoptosis. In contrast, the apoptotic regulatory protein interacting with LARP6 has not yet been identified in mammals. Further investigations are therefore needed to determine whether LARP6 exerts its anti-apoptotic effects through interaction with analogous apoptotic inhibitory proteins. Despite this gap in knowledge, multiple studies in cancer cells and fibroblasts have demonstrated that LARP6 promotes cell proliferation, enhances survival, and facilitates invasion [37, 39, 40]. Collectively, these findings suggest that LARP6 may possess a conserved pro-survival function across diverse cell types, potentially extending to vascular smooth muscle cells in the context of atherosclerosis.

Our analysis of human bulk RNA-seq data revealed a significant decrease in LARP6 within vulnerable atherosclerotic plaques. Pathway enrichment analysis showed that genes co-regulated with LARP6 were predominantly associated with ECM organization, suggesting that LARP6 may contribute to plaque stability by regulating ECM integrity. Moreover, we demonstrated that LARP6 protein expression is negatively regulated by miR-1976, a microRNA previously reported by Cai et al. to be markedly upregulated in patients with cardiovascular disease compared with healthy individuals [41]. The inverse relationship between miR-1976 and LARP6 provides a potential mechanistic link between microRNA dysregulation and compromised ECM stability in vulnerable plaques.

Taken together, our findings identify LARP6 as a previously unrecognized regulator of plaque stability and, to our knowledge, represent the first description of LARP6 function in the context of atherosclerosis. These results highlight LARP6 as a potential therapeutic target for reinforcing fibrous cap integrity and preventing plaque rupture.

## Acknowledgement

M. Zhang participated in the study design, collected and analyzed data, interpreted results, drafted the manuscript, and prepared figures. S. Danchuk collected data and provided technical expertise. S. Sukhanov assisted with experimental design and data interpretation. T. Yoshida provided key technical support and contributed to data interpretation. P. Delafontaine conceived and designed the study and revised the manuscript. Y. Higashi conceived and designed the research, interpretated data, and revised the manuscript before final approval. All authors read and approved of the final manuscript. All authors acknowledge extensive support and technical contributions by Zhaohui Li (University of Missouri School of Medicine).

---

**Table S1.**
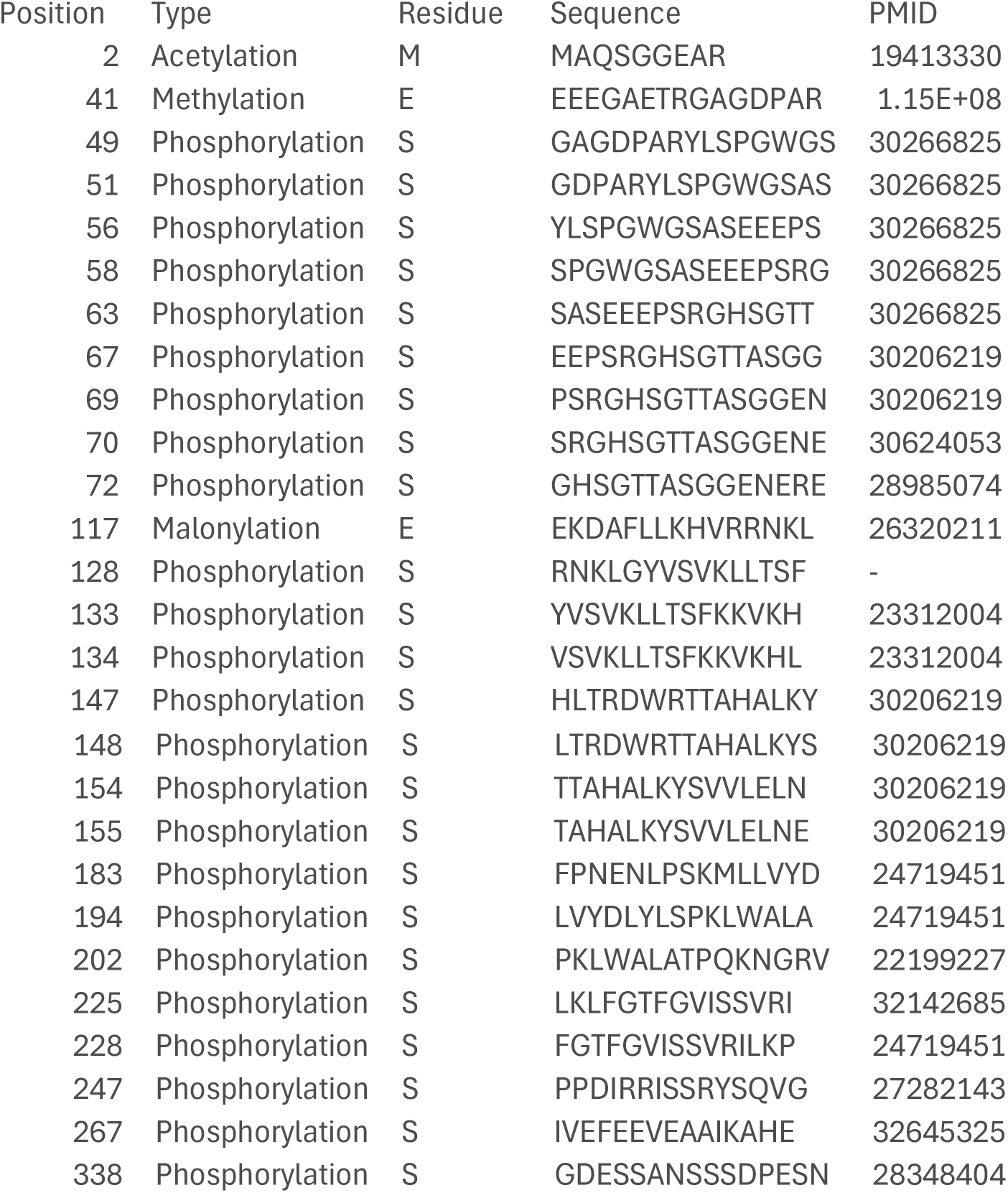
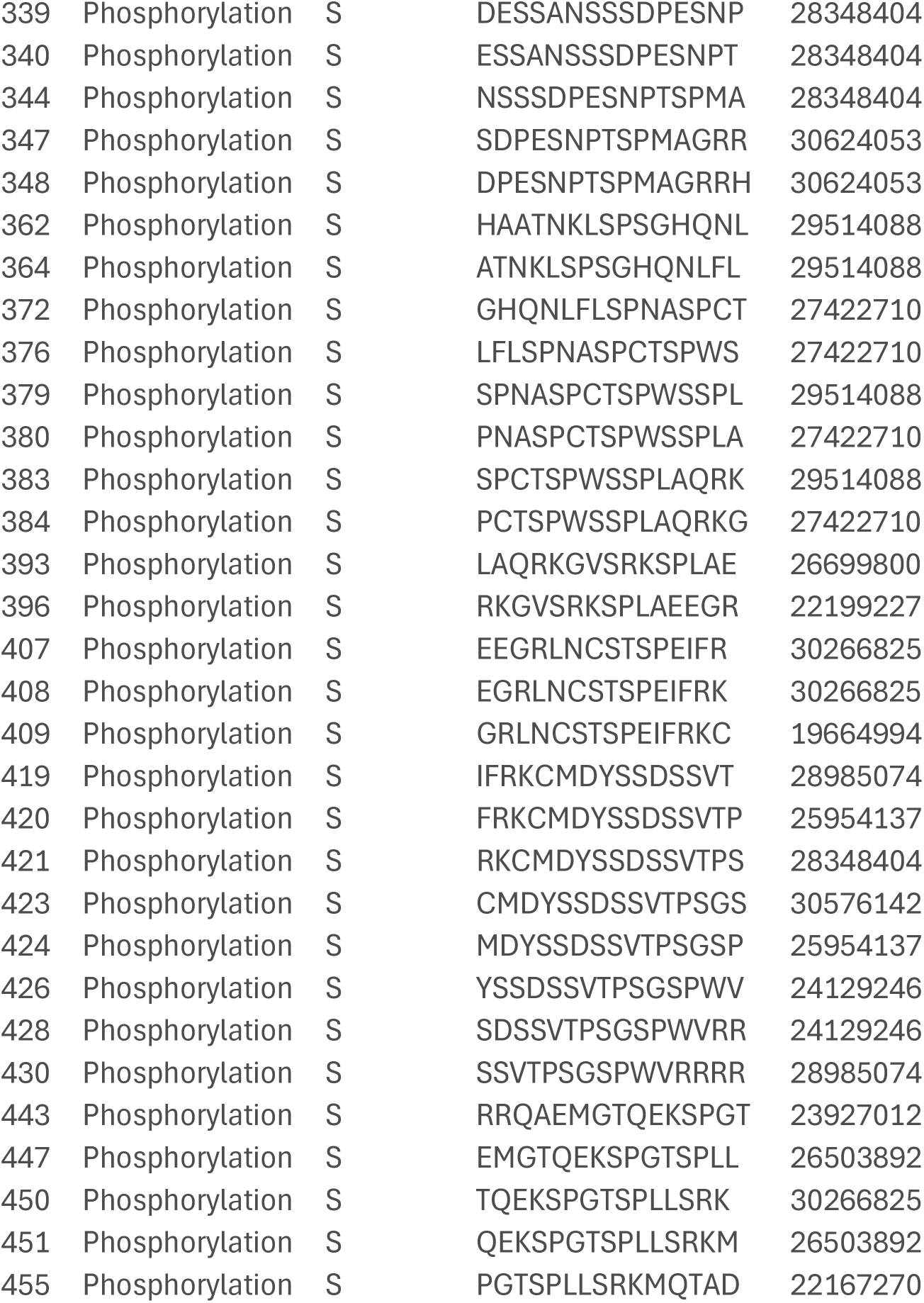
Post translational modification sites on LARP6.

**Table S2.** miRs in hAoSMCs which are regulated by IGF-1.

| miRs | Relative expression | P value |  |
| --- | --- | --- | --- |
| miR-126-3p | 0.75±0.01 | ** |  |
| miR-126-5p | 0.82±0.03 | NS |  |
| miR-182-5p | 0.80±0.10 | NS |  |
| miR-495 | 0.93±0.05 | NS |  |
| miR-744-5p | 0.76±0.00 | *** |  |
| miR-1976 | 0.59±0.07 | *** | **P<0.01, ***P<0.001 |

**Fig. S1.**
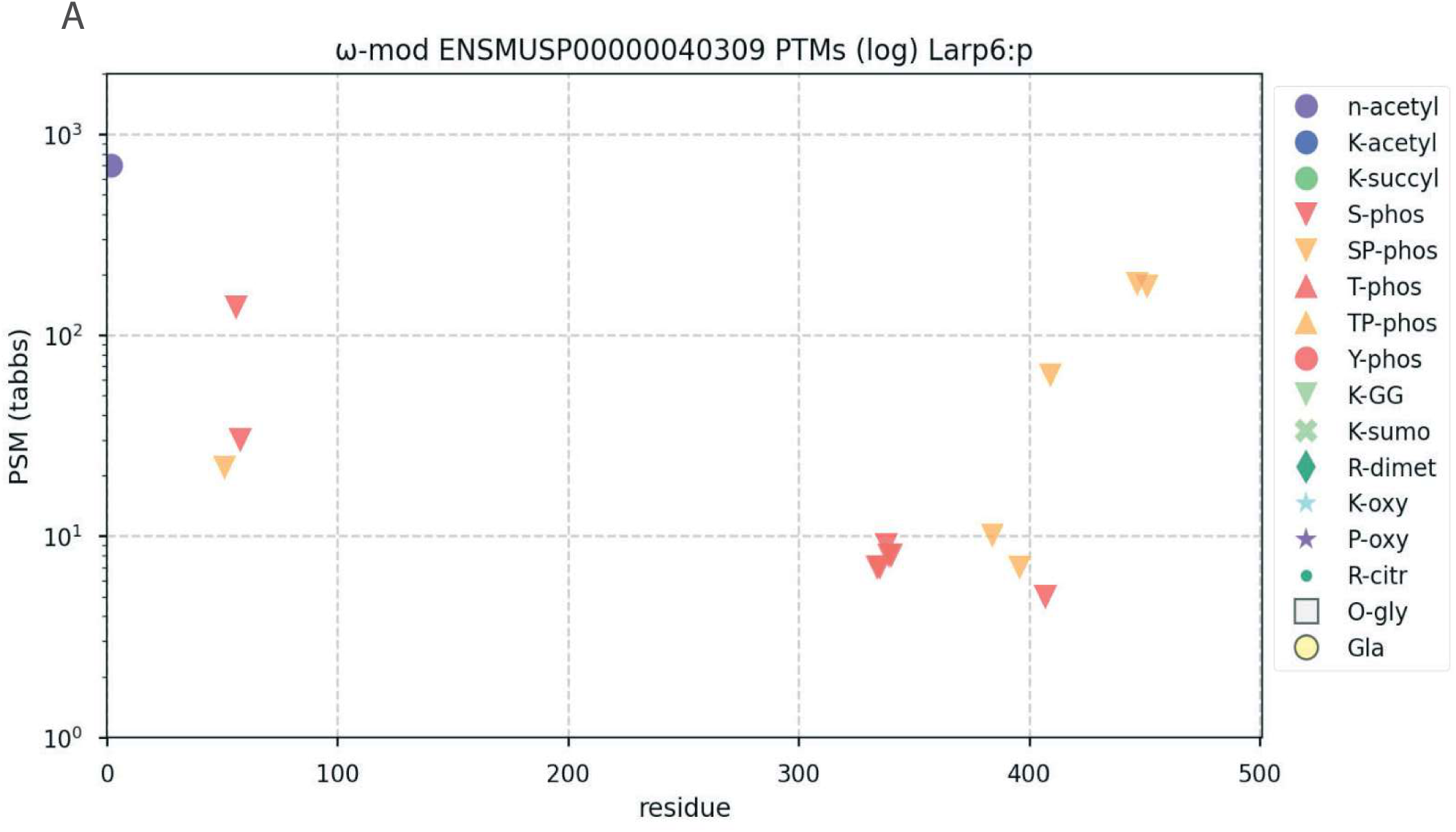
Post-translational modifications identified in mouse Larp6 (ENSMUSP00000040309). Post-translational modification (PTM) sites detected by mass spectrometry are plotted according to their amino acid residue position (x-axis) and peptide spectral match (PSM) abundance on a log scale (y-axis). Each symbol denotes a specific modification type, including acetylation, succinylation, phosphorylation (S/T/Y), ubiquitin-like K-GG, SUMOylation, dimethylation, oxidation, citrullination, and O-glycosylation, as indicated in the legend. The distribution highlights multiple phosphorylation-enriched regions and diverse PTMs across both N-terminal and C-terminal domains of Larp6.

**Figure S2.**
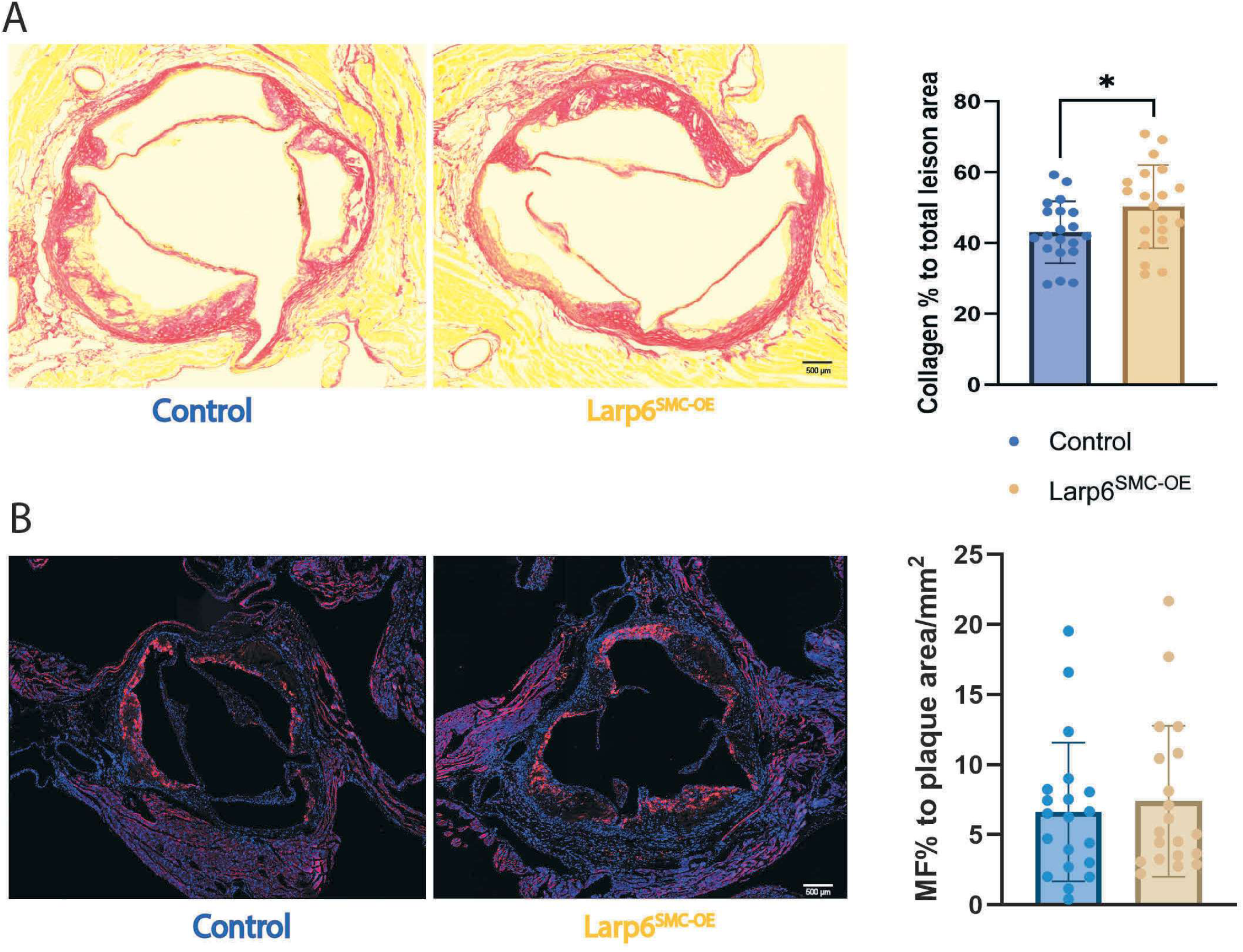
Larp6 overexpression in SMCs increases collagen content without altering MF-like SMC abundance in atherosclerotic lesions. **A,** Representative Picrosirius Red-stained aortic root sections from Control and Larp6SMc-oE mice. Quantification of collagen area expressed as percentage of total lesion area shows a significant increase in collagen content in Larp6SMc-oE plaques (right). *n* = 20 per group. Scale bar: 500 μm. **B,** Representative immunofluorescence images showing MF-like SMCs (red) within the aortic root lesions of Control and Larp6SMc-oE mice. Quantification of MF-like SMC area normalized to plaque area indicates no significant difference between groups (right). *n* = 20 per group. Scale bar: 500 μm. Data are presented as mean± SEM. *P<* 0.05 was considered significant.

**Figure S3.**
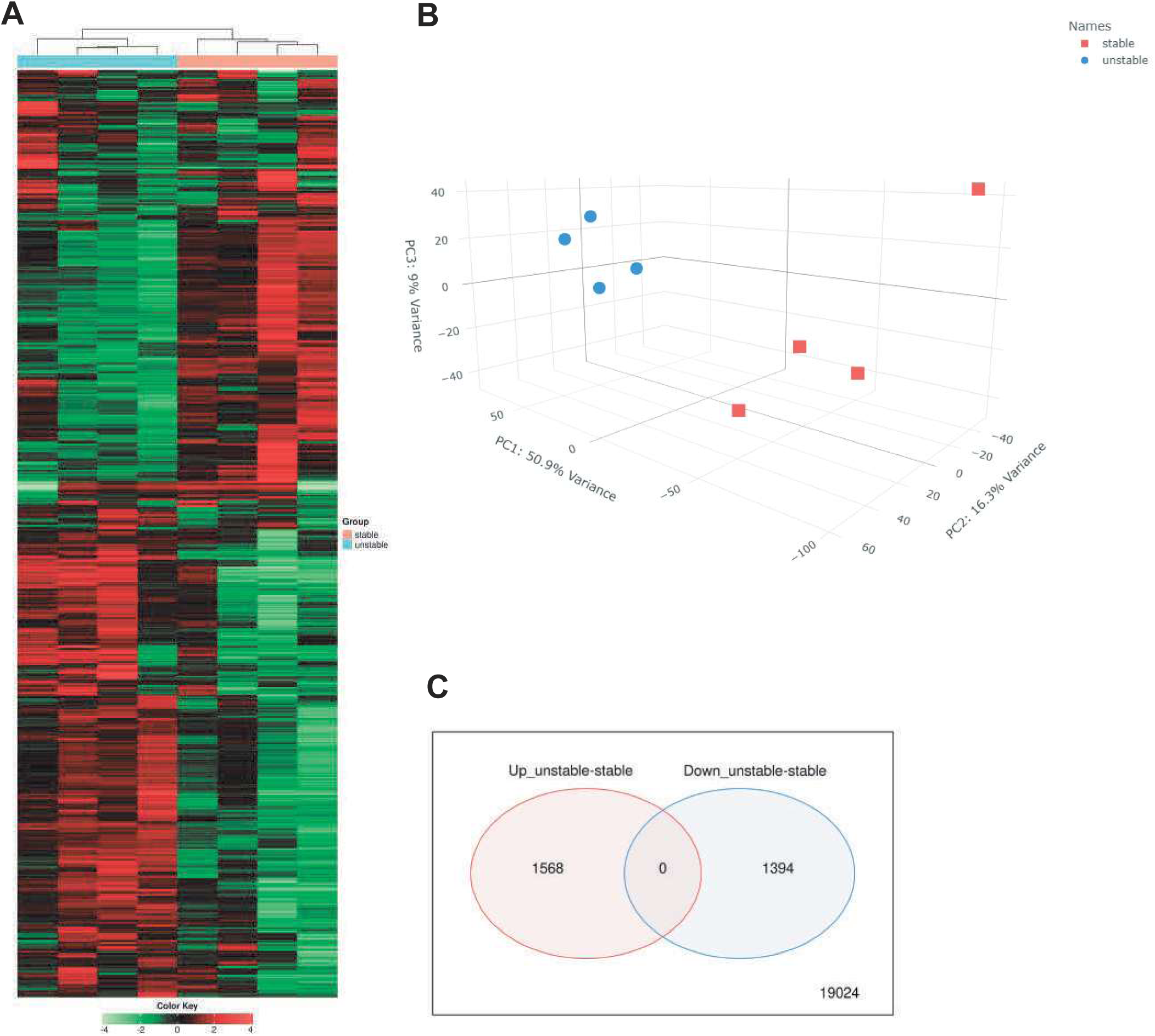
Transcriptomic differences between stable and unstable atherosclerotic plaques. **A,** Unsupervised hierarchical clustering heatmap of differentially expressed genes (DEGs) in stable vs. unstable lesions. Each column represents an individual sample, and rows represent genes scaled by Z-score. Distinct clustering patterns indicate strong transcriptional separation between plaque phenotypes. **B,** Principal component analysis (PCA) illustrating global transcriptomic variance across samples. Stable plaques (red) and unstable plaques (blue) segregate along PC1 and PC2, accounting for the majority of expression variance and demonstrating clear sample-level grouping. **C,** Venn diagram showing the number of genes significantly upregulated and downregulated in unstable vs. stable plaques. A total of 1,568 genes were upregulated and 1,394 genes were downregulated, with no overlap between the two sets under the applied thresholds.

